# Direct binding of the flexible C-terminal segment of periaxin to β4 integrin suggests a molecular basis for CMT4F

**DOI:** 10.1101/524793

**Authors:** Arne Raasakka, Helen Linxweiler, Peter J. Brophy, Diane L. Sherman, Petri Kursula

## Abstract

The process of myelination in the nervous system requires coordinated formation of both transient and stable supramolecular complexes. Myelin-specific proteins play key roles in these assemblies, which may link membranes to each other or connect the myelinating cell cytoskeleton to the extracellular matrix. The myelin protein periaxin is known to play an important role in linking the Schwann cell cytoskeleton to the basal lamina through membrane receptors, such as the dystroglycan complex. Mutations that truncate periaxin from the C terminus cause demyelinating peripheral neuropathy, Charcot-Marie-Tooth disease type 4F, indicating a function for the periaxin C-terminal region in myelination. We identified the cytoplasmic domain of β4 integrin as a specific high-affinity binding partner for periaxin. The C-terminal region of periaxin remains unfolded and flexible when bound to the third fibronectin type III domain of β4 integrin. Our data suggest that periaxin is able to link the Schwann cell cytoplasm to the basal lamina through a two-pronged interaction *via* different membrane protein complexes, which bind close to the N and C terminus of this elongated, flexible molecule.

## Introduction

Long axonal segments in the vertebrate peripheral nervous system (PNS) are ensheathed by myelin, which accelerates nerve impulse propagation and which supports axons both mechanically and trophically (1). A Schwann cell wraps its plasma membrane, partially excluding its cytosol, several times around a selected axonal process. This results in compact myelin, which has high lipid and protein content and is responsible for axonal insulation. Compact myelin is surrounded by a narrow compartment with higher cytosolic content, non-compact myelin, which acts as a supportive metabolic compartment to ensure long-term myelin stability (2). Both myelin compartments contain a specific selection of proteins that have distinct tasks in ensuring the correct formation and stability of myelin: failure may result in one of several disease states, including the peripheral inherited neuropathies Charcot-Marie-Tooth (CMT) disease and Dejerine-Sottas syndrome (DSS). A large number mutations in different PNS proteins has been linked to these conditions (3,4). On the other hand, only a handful of CMT mutations have been characterized at the molecular structural level in order to understand the fine details of disease mechanisms (5–7).

The formation of myelin in the CNS and PNS, as well as its lifelong maintenance, requires an intricate network of molecular interactions that link the myelin membrane, the cytoskeleton of the myelinating glial cell, and the extracellular matrix or the axonal surface together into a large supramolecular complex. A number of proteins, many of which seem specific for myelinating cells, have been pinpointed as playing roles in myelination; however, often the molecular details of the relevant processes and protein-protein interactions remain unknown. Myelin proteins have been specifically highlighted as a knowledge gap in structural biology in the past (8), although more structural data from myelin proteins are becoming available. However, structures of protein-protein complexes of myelin-specific proteins mainly remain uncharacterized to date.

Unlike oligodendrocytic myelin in the CNS, Schwann cells in the PNS are surrounded by a carbohydrate-rich basal lamina, which is adhered to the outermost (abaxonal) Schwann cell plasma membrane bilayer *via* dystroglycans and α6β4 integrin, contributing to the mechanical stability of myelinated nerves (9–11). Additionally, non-compact PNS myelin contains tight membrane-apposed structures at the abaxonal layer. These structures surround cytosolic channels of non-compact myelin, referred to as Cajal bands, which contain substantial microtubule-based transport as well as ribosomal activity (12,13). The membrane appositions have tight morphology and are enriched in periaxin (PRX) – the most abundant PNS non-compact myelin protein (14). Cajal bands and the membrane appositions are important in regulating myelin stability. Furthermore, PRX influences the myelin sheath internode distance, and thus influences nerve conduction velocity (15,16). These structures can be disturbed in human demyelinating diseases, as well as in corresponding mouse models (16–19).

Two isoforms of PRX are generated through alternative splicing (20). Disease mutations in PRX often truncate the long C-terminal region of the larger L-PRX isoform (21). The molecular mechanism of disease in these cases has remained enigmatic, as the best-characterized protein interactions and functions of L-PRX so far lie very close to the N terminus. These include the PDZ-like domain, which mediates homo- and heterodimerization of PRX (18,22,23), and the segment after it, which is known to bind dystrophin-related protein 2 (DRP2) and link PRX to the dystroglycan complex (18). A conceivable additional mechanism of PRX function and involvement in disease could involve specific protein interaction sites at the C-terminal, isoform-specific end of L-PRX.

We wanted to identify novel binding partners for the L-PRX C-terminal region. The third cytoplasmic fibronectin-type III (FNIII) domain of β4 integrin (β4-FNIII-3) was identified as a high-affinity binder, and the complex was characterized using biophysical and structural biology techniques. The observed direct molecular interaction is likely to be important for the function of PRX and β4 integrin in developing and mature myelin, and it provides a molecular basis for PRX mutations in CMT that result in the expression of truncated L-PRX.

## Materials and methods

### Yeast two-hybrid screening

Yeast two-hybrid screening was performed essentially as previously described (18). Briefly, a random-primed rat sciatic nerve cDNA library in λACTII was screened using the C-terminal region of rat L-PRX (residues 681-1383) in the pAS2-1 vector (Clontech) as bait. Three independent clones of β4 integrin each containing the third FNIII domain were found. To identify the domain of PRX, which interacted with β4 integrin, deletion contructs of PRX vwere made by PCR and subcloned into pAS2-1. β-galactosidase activity was tested by filter lift assays with one of the β4 integrin clones (62BpACTII).

### GST pulldown

The β4 integrin FNIII-3 domain (amino acids 1512-1593) cDNA was amplified by PCR and subcloned into pGEXKG. As a control, the adjacent fourth FNIII domain was cloned into the same vector. The recombinant protein was expressed and purified using a Glutathione-Sepharose 4B column as described (18). GST pulldowns were performed by incubation of the GST fusion protein bound to Glutathione-Sepharose with a sciatic nerve lysate, as previously described (18).

### Immunoaffinity chromatography

Immunoaffinity pull-downs were performed essentially as described (18). PRX and β4 integrin in the pull-down fractions were identified by Western blot. PRX antibodies have been described (24). Antibodies against β4 integrin were raised in rabbits using a peptide corresponding to amino acids 1756-1772, to which an N-terminal cysteine was attached for coupling to keyhole limpet hemocyanin (CTEPFLIDGLTLGTQRLE), as described (20).

### Immunofluorescence

Mice were perfused intravascularly with 4% paraformaldehyde in 0.1 M sodium phosphate buffer (pH 7.3) and sciatic nerve cryosections were prepared and immunostained as described (25). Antibodies against PRX have been described (26), and the monoclonal antibody against β4 integrin was a generous gift from Dr. S.J. Kennel, Biology Division, Oak Ridge National Laboratory.

### Transfection and coimmunoprecipitation

Full-length cDNA clones encoding rat β4 integrin and human α⍰ integrin were subcloned into the expression vectors pcDNA3.1 and pRC/CMV, respectively, and were generous gifts from Dr M.L. Feltri, Hunter James Kelly Research Institute, University of Buffalo and Dr Arnoud Sonnenberg, Netherlands Cancer Institute. PRX cDNA in the expression vector pCB6 has been described (27). PRX, β4 integrin, and α⍰ integrin were expressed in HEK293 cells. After transfection, the proteins were immunoprecipitated as described (18).

### Proteomics

Proteomics analyses were carried out exactly as described (24), comparing the PRX-bound proteomes of PRX^-/-^ mice carrying a transgene for either wild-type PRX (PrxTg/PRX^-/-^) or PRX with a C-terminal truncation mutation (mouse L-PRX truncated at residue 1016; ΔCPrxTg/PRX^-/-^) corresponding to human CMT4F R1070X (28,29). Proteins from mouse nerve lysates were crosslinked followed by immunoprecipitation with a PRX antibody, and then subjected to mass spectrometry (MS) analyses. Normalized abundance of β4 integrin was determined by MS using Progenesis (Nonlinear Dynamics) as described (24).

### Recombinant protein production

Synthetic genes encoding the rat L-PRX C-terminal region (UniProt ID: Q63425, amino acids 1036 – 1383; PRX-C) and rat β4-FNIII-3 (UniProt ID: Q64632, amino acids 1512 – 1602) were ordered from DNA2.0 (Newark, CA, USA) in the pJ401 bacterial expression vector. An additional sequence encoding for an N-terminal hexahistidine tag, a short linker, and a tobacco etch virus (TEV) protease digestion site (MHHHHHHSSGVDLGTENLYFQS) were included at the start of the protein-coding insert.

PRX-C was expressed in *E. coli* BL21(DE3) using 0.4 mM IPTG induction for 1.5 h in LB medium containing 100 μg/ml ampicillin, at 37 °C. After expression, the cells were pelleted by centrifugation and sonicated in Ni-NTA washing buffer (40 mM HEPES, 400 mM NaCl, 6 M urea, 20 mM imidazole, 1 mM PMSF, pH 7.5) supplemented with protease inhibitors (Roche). Purification was performed using Ni-NTA affinity chromatography and standard procedures. Elution was done with 32 mM HEPES, 320 mM NaCl, 4.8 M urea, 500 mM imidazole, pH 7.5. The eluted fraction was dialyzed at 4 °C with constant stirring against 40 mM HEPES, 400 mM NaCl, 1 mM DTT, pH 7.5, before addition of recombinant TEV protease for affinity tag removal (30). The digestion was allowed to proceed overnight while dialyzing, which resulted in cleaved PRX-C with an additional N-terminal Ser residue. The protein was subjected to a 2^nd^ Ni-NTA step, in the absence of urea. The unbound and wash fractions were combined and concentrated, and sequential size exclusion chromatography (SEC) on a HiLoad Superdex 200 pg 16/600 column (GE Healthcare) was used to separate the cleaved protein from contaminants and degradation products. Depending on downstream application, either 20 mM HEPES, 300 mM NaCl, 1% (v/v) glycerol, 0.5 mM TCEP, pH 7.5 (SEC buffer) or 20 mM HEPES, 150 mM NaCl, pH 7.5 (HBS) was used as running buffer.

β4-FNIII-3 was expressed in *E. coli* BL21(DE3) in LB or TB medium containing 100 μg/ml ampicillin, with 0.4 mM IPTG induction for 3 h at 37 °C. The cells were harvested as above and resuspended in 40 mM HEPES, 400 mM NaCl, 20 mM imidazole, pH 7.5. After lysis by sonication, a Ni-NTA chromatography was carried out essentially as described above, omitting urea in all buffers. After Ni-NTA, the eluted protein was directly subjected to SEC using a HiLoad Superdex 75 pg 16/60 column (GE Healthcare) using either HBS or SEC buffer.

### Pulldown experiment with purified proteins

Purified recombinant β4-FNIII-3 and PRX-C were used in pulldown experiments to confirm the direct interaction. PRX-C with and without the His-tagged β4-FNIII-3 was mixed with Ni-NTA agarose for 1 h at +4 °C, in binding buffer (20 mM HEPES, 150 mM NaCl, 20 mM imidazole, pH 7.5). The matrix was collected by centrifugation at 150 g for 5 min at +4 °C. Three washes with binding buffer were carried out, and proteins were eluted with 20 mM HEPES, 150 mM NaCl, 500 mM imidazole, pH 7.5. All fractions were analyzed with SDS-PAGE.

In addition, a partially degraded PRX-C sample was employed to identify fragments that do and do not bind β4-FNIII-3. The pulldown experiment was carried out exactly as above. Bands were excised from the gel and processed for tryptic peptide mapping.

### Synchrotron radiation circular dichroism (SRCD) spectroscopy

SRCD data were collected from 0.15 – 0.6 mg ml^-1^ protein samples in 20 mM Na-phosphate, 150 mM NaF, pH 7.5 on the AU-CD beamline at the ASTRID2 synchrotron (ISA, Aarhus, Denmark). 100-μm pathlength closed circular cells (Suprasil, Hellma Analytics) were used for the measurements. Spectra were recorded from 170 to 280 nm at 20 °C. Temperature scans were performed from 10 to 90 °C in 5 °C intervals, with 5 min incubation per time point prior to spectral acquisition.

Buffer spectra were subtracted from the protein samples, and CD units were converted to Δɛ (M^-1^ cm^-1^), using rPRX-C concentration determined using refractometry and/or rFNIII-3 concentration determined using absorbance at 280 nm. Deconvolution was performed using DichroWeb (31) with the CDSSTR algorithm (32) and SP175 reference set (33), or using BeStSel (34). Secondary structure prediction was performed using JPred (35).

### Small-angle X-ray scattering (SAXS)

Synchrotron SEC-SAXS data for PRX-C, β4-FNIII-3, and their complex were collected on the B21 beamline at Diamond Light Source (Didcot, UK) using an on-line size exclusion setup: the chromatography was performed using an Agilent 1200-series HPLC system and a Superdex 200 increase 3.2/300GL (GE Healthcare) column with 20 mM Tris-HCl, 150 mM NaCl, pH 7.4 as mobile phase at an isocratic flow of 0.04 ml/min. 45 μl injections of 6.5 – 9.8 mg/ml total protein were performed for PRX-C, β4-FNIII-3, and their equimolar complex (145 μM each). Scattering was directly recorded from the eluted proteins at 6 s exposure per frame, 591 frames per run. The frames containing a stable R_g_ within an eluted *I*_0_ peak were selected and combined using ScÅtter (http://www.bioisis.net/tutorial). Data were processed and analyzed using the ATSAS package (36). GNOM was used to calculate distance distribution functions (37), and *ab initio* modeling was performed using GASBOR (38). Multiphase modeling of protein complex data was performed using MONSA (39) and ensemble optimization analysis with EOM (40). See Supplementary Table 1 for further details.

For IDPs, more accurate values of R_g_ can be obtained from SAXS data using the Debye formalism. Briefly, by plotting [I(s)]^-1^ *vs*. s^2.206^, in the range (s R_g_)^2^<3, one can use much more data than in a regular Guinier plot. Here, a linear fit of the plot is used to obtain R_g_ = (a/0.359b)^0.453^; a = slope, b = y intercept (41). Theoretical values for R_g_ and D_max_ for a random chain can be calculated from sequence length as follows: R_g_ = R_0_N^v^; R_0_ = 1.98 Å, N = number of residues, v = 0.602 (42) for denatured proteins; R_0_ = 2.54 Å, v = 0.522 (43) for IDPs. Average end-to-end distance can be estimated as <L^2^> = L_0_N; L_0_ = 81.8 Å (42).

### Mass spectrometry and covalent crosslinking

The molecular weight of PRX-C and β4-FNIII-3 was verified by mass spectrometry. In short, the undigested masses were determined using ultra-performance liquid chromatography (UPLC) coupled electrospray ionization (ESI) time-of-flight mass spectrometry in positive ion mode using a Waters Acquity UPLC-coupled Synapt G2 mass analyzer with a Z-Spray ESI source.

Protein crosslinking was carried out to conjugate surface-exposed carboxylate sidechains with lysines with a zero-length crosslinker. All crosslinking reactions were carried out at 40 μM final protein concentration in 100 mM bis-tris methane, 150 mM NaCl, 4 mM 1-ethyl-3-(3-dimethylaminopropyl)carbodiimide, 20 mM *N*-hydroxysuccinimide, pH 6.5. The activation step was allowed to proceed for 15 min at ambient temperature, followed by quenching the reactions through adding 2-mercaptoethanol to 20 mM. After addition of the second protein in selected reactions, incubation was continued for another 3 h at ambient temperature and stopped by adding ethanolamine to 10 mM. The reactions were analyzed using SDS-PAGE.

The crosslinked proteins were identified using matrix-assisted laser desorption/ionization time-of-flight (MALDI-TOF) mass spectrometry. From stained SDS-PAGE gels, gel bands were cut, staining removed by sequential washing with 50 mM NH_4_HCO_3_ in 40% acetonitrile (ACN). Proteins were subjected to in-gel Cys reduction using 20 mM DTT and subsequent alkylation using 40 mM α-iodoacetamide. After this, all proteins were in-gel digested (20 ng of trypsin (Sigma-Aldrich) per gel piece), followed by peptide extraction from gel pieces using 30% ACN/0.1% trifluoroacetic acid (TFA), and transfer to a Bruker anchor plate. 800 μg/ml α-cyano-4-hydroxy cinnamic acid in 85% ACN/0.1% TFA with 1 mM NH_4_H_2_PO_4_ was used as matrix. Peptide mass spectra and MS/MS spectra were measured with a Bruker Ultra fleXtreme MALDI-TOF mass spectrometer and compared to theoretical spectra generated using the known protein sequences.

### Multi-angle static and quasielastic light scattering

SEC-MALS and quasielastic light scattering (QELS) data were collected to determine the monodispersity, hydrodynamic radius, and molecular weight of PRX-C, β4-FNIII-3, and their equimolar complex (145 μM each). The chromatography was performed using an Agilent 1200-series HPLC system and a Superdex 200 increase 3.2/300GL (GE Healthcare) column with 20 mM Tris, 150 mM NaCl, pH 7.4 as mobile phase. Protein samples of 160 – 250 μg were injected into the column at an isocratic flow of 0.04 ml/min, and light scattering was recorded using a Wyatt miniDAWN HELEOS-II instrument with 18 detectors and a QELS module at ambient temperature. The refractive index was measured using a Wyatt Optilab T-rEX refractometer and used as the concentration source. All data were analyzed using the ASTRA software (Wyatt).

### Protein crystallography

β4-FNIII-3 was crystallized using sitting-drop vapor diffusion in drops consisting of 150 nl protein solution (12.7 mg/ml in SEC buffer) mixed with 150 nl of reservoir solution. Initially, crystals formed in a wide variety of PEG-based conditions in PACT Premier and JCSG+ (Molecular Dimensions) crystal screens. Optimized crystals used for diffraction data collection were grown at 20 °C in conditions containing 16 – 22% (w/v) PEG 3350 and 180 – 240 mM NH_4_NO_3_.

Prior to diffraction data collection, the crystals were cryoprotected briefly by adding 1.5 μl of cryoprotectant solution directly into the sitting drop mix. The cryoprotectant consisted of 75% (v/v) well reservoir and 25% (v/v) PEG 400. After soaking in cryoprotectant, crystals were mounted in loops and snap-frozen in liquid N_2_.

X-ray diffraction data were collected at 100 K on the P13 beamline, EMBL/DESY, Hamburg, Germany (44) and the ID30A-1 beamline at ESRF (Grenoble, France) (45,46). Data were processed using XDS (47). Phasing was performed with molecular replacement using the human β4-FNIII-3 structure (PDB ID 4wtw) (48) as the search model in Phaser (49). The structure was refined in phenix.refine (50), and model building was done in Coot (51). The low solvent content of 36% (including ordered water) leads to a higher-than-average difference between R_work_ and R_free_ during structure refinement.

The structure was validated and analyzed using DSSP (52), MolProbity (53), PyMOL, PDB2PQR (54), APBS (55), and UCSF Chimera (56). The crystal structure was subjected to atomistic molecular dynamics simulations for 550 ns in YASARA (57), essentially as described (58).

### Isothermal titration calorimetry (ITC)

PRX-C and β4-FNIII-3 were dialyzed into HBS overnight to ensure matching buffer. The proteins were passed through a 0.22-μm filter, and concentrations were checked. ITC was performed at 25 °C using a Malvern MicroCal iTC200 calorimeter with reference power set to 5 μcal/s. 680 μM β4-FNIII-3 was titrated into 350 μl of μM PRX-C under constant stirring. A total of 38 1-μl injections were performed, with a 240-s waiting period between injections. Data were analyzed using Origin. The titration experiment was repeated twice with two separate protein batches and nearly identical values were obtained in each case.

### Thermal stability assays

Thermal stability experiments were performed for 0.25 – 2 mg/ml β4-FNIII-3 and a μM equimolar complex of PRX-C and β4-FNIII-3 in SEC buffer using a label-free fluorescence-based approach (nanoDSF). The instrument used was a NanoTemper Prometheus NT.48 nanoDSF with a backscattering option to detect aggregation onset. Each 10-μl sample was loaded inside a glass capillary, and a constantly monitored scan from 20 – 95 °C using a 2 °C/min ramp rate was performed. Fluorescence at emission wavelengths 330 nm and 350 nm (F_330_ and F_350_, respectively) was monitored, and a single transition event was observed in the fluorescence ratio (F_350_/F_330_) in all samples. Melting temperature midpoint (*T*_m_) values were extracted from the 1^st^ derivative peak maximum.

## Results

### Identification of the cytoplasmic domain of β4 integrin as a potential periaxin ligand

In order to identify putative protein ligands for the C-terminal segment of L-PRX, which is missing in the presence of the CMT4F R1070X mutation (24,28), we carried out a yeast 2-hybrid screen of a rat sciatic nerve cDNA library with the C-terminal segment, missing upon the R1070X mutation, as bait. As a result, we obtained three separate clones containing the β4-FNIII-3 domain (Figure 1A). One of these (clone 62B) was further used in confirmatory experiments.

**Figure 1.**
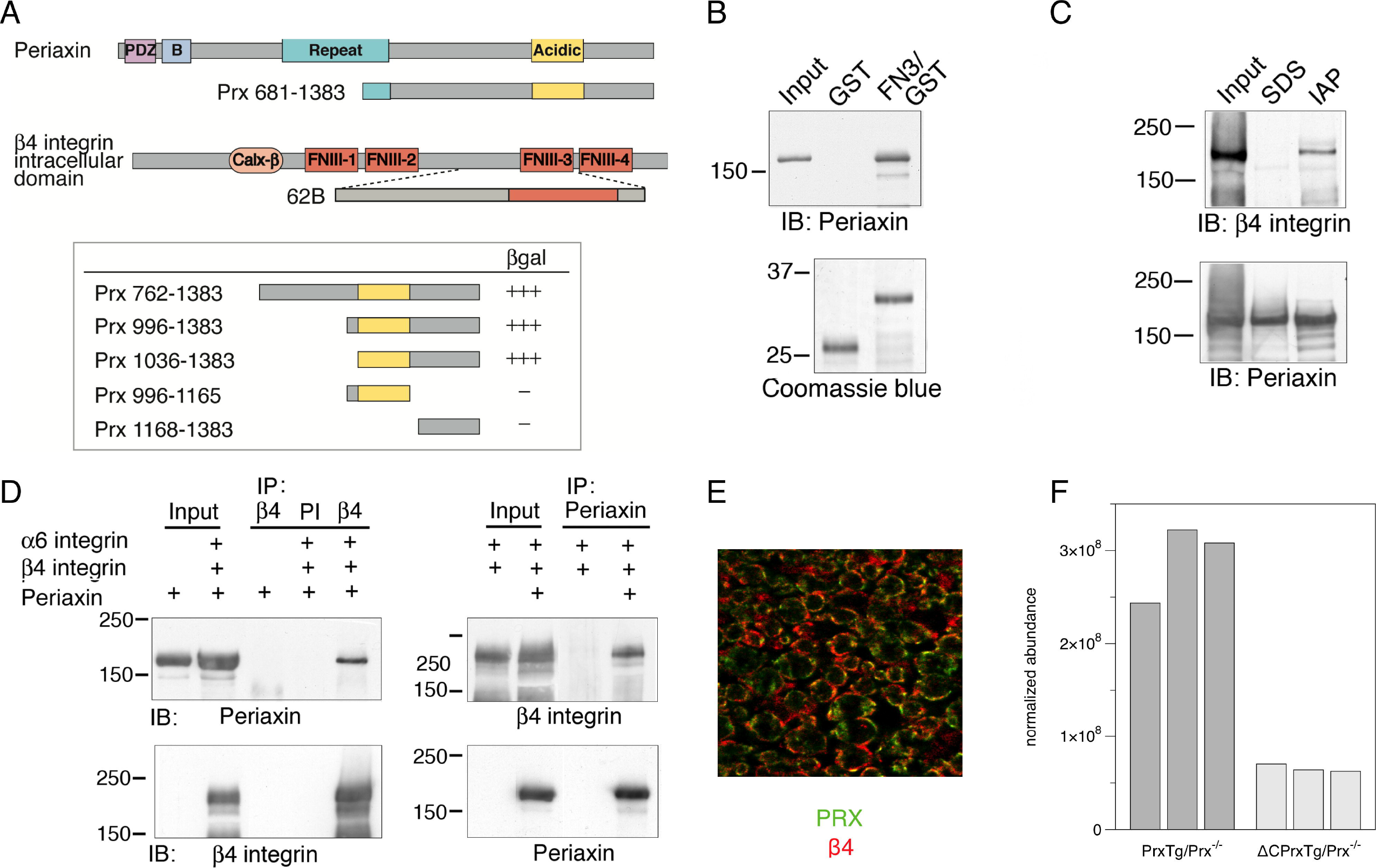
Identification of an interaction between periaxin and β4 integrin. A. Top: structure of full-length L-PRX and the bait used in a yeast two-hybrid screen using the C terminus of L-PRX (aa 681-1383) fused to the GAL4 DNA-binding domain. Bottom: structure of the intracellular domain of β4 integrin and one (62B) of the three clones recovered from the screen, all of which included the third FNIII domain of β4 integrin. B. Identification of the region of PRX, which binds to β4 integrin. The strength of interaction between the 62B clone and a series of PRX constructs in a yeast two-hybrid β-galactosidase assay was assessed semiquantitatively by the time taken for colonies to turn blue (+++, 30 min; ++, 30–60 min; +, 60–180 min). C. Interaction of β4-FNIII-3 with PRX *in vitro*. GST or a GST-β4-FNIII-3 fusion protein were incubated with a sciatic nerve lysate *in vitro* and any bound L-PRX was detected by Western blotting (IB) using a PRX antibody after SDS-PAGE. The input lane confirms the presence of PRX in the lysate. Coomassie blue staining shows that equivalent amounts of GST and the GST fusion proteins were used. D. Immunoaffinity copurification of β4 integrin and PRX. Detergent extracts of mouse sciatic nerve in the non-ionic detergent Igepal were incubated with beads to which affinity-purified sheep anti-PRX antibodies had been covalently coupled. After extensive washing, bound proteins were eluted and analyzed by SDS-PAGE and Western blotting. The SDS control lane contains proteins that bound to the beads after first solubilizing the nerves in SDS to disrupt protein-protein interactions, followed by dilution of the SDS with Triton X-100. E. Coimmunoprecipitation of β4 integrin and PRX from transfected HEK293 cells. PRX and β4 integrin were detected by Western blot (IB) after immunoprecipitation (IP) with β4 integrin antibodies when β4 integrin, α6 integrin, and PRX were coexpressed. Preimmune serum (PI) did not precipitate either protein. Reciprocally, β4 integrin and PRX were detected by Western blot after immunoprecipitation (IP) with anti-PRX antibodies, when β4 integrin, α6 integrin, and PRX were coexpressed. F. Immunohistochemistry. Both PRX (green) and β4 integrin (red) localize at the abaxonal membrane in mature myelin. G. Quantification of β4 integrin from the immunoprecipitation of crosslinked sciatic nerves from both the full-length PRX transgenic mouse on a PRX null background (n=3) versus C-terminally truncated PRX transgene on a PRX null background (n=3).

The interaction was confirmed and the binding site coarsely mapped by using different domains of L-PRX in the screen, with the obtained β4 integrin construct 62B as bait (Figure 1A). The results indicate that the last ∼350 residues, containing the acidic region of L-PRX, are crucial for the interaction in this system. Interestingly, both PRX(996-1165) and PRX(1168-1383) gave negative results, suggesting that the binding site may be located close to the point separating these two constructs, or that the site might be a combination of more than one segment required for high-affinity binding.

### Evidence for β4 integrin-PRX interaction in tissues and cells

As the next step, GST pulldowns from sciatic nerve lysates were carried out using a GST-β4-FNIII-3 construct. Western blotting identified L-PRX in the fraction bound to the GST fusion protein, but not GST alone (Figure 1B). DRP2 was also identified in the fraction pulled down by GST-β4-FNIII-3, while GST-β4-FNIII-4 was unable to pull down PRX or DRP2 (Supplementary Figure 1). Immunoaffinity pulldowns were further carried out using an anti-PRX antibody, and an analysis of the fractions showed copurification of L-PRX and β4 integrin (Figure 1C), when SDS - breaking up protein complexes - was not used during extraction.

The putative interaction between L-PRX and β4 integrin was further studied using co-immunoprecipitation from cultured HEK293 cells overexpressing both proteins. After immunoprecipitation with either β4 integrin or PRX antibodies, both proteins were observed in the eluted fraction, showing they are present in the same complex (Figure 1D).

To observe the localization of PRX and β4 integrin in myelinating glial cells, immunofluorescence staining of mouse sciatic nerves was carried out. The result confirms earlier studies (9,59,60), in that both PRX and β4 integrin are localized at the outermost membrane layer of myelin (Figure 1E).

### Changes in the L-PRX interactome in a mouse model of CMT4F

Using the mouse model for CMT4F (24), which carries a truncating mutation mimicking human L-PRX with the R1070X mutation, we carried out proteomics experiments to follow the expression level of possible PRX ligand proteins in this model. PRX was immunoprecipitated from mouse nerves after crosslinking, and the bound proteins were analyzed. The levels of DRP2 were decreased in the PRX interactome of these mutant animals (24). In addition, a several-fold drop in the level of β4 integrin could here be observed in the mutant compared to the wild-type animals based on data from the same experiment (Figure 1F). The result provides strong evidence for a functional protein complex involving L-PRX and β4 integrin in mouse peripheral nerve myelin *in vivo*.

### Complex of purified recombinant β4 integrin and PRX

The above experiments provided evidence for a direct interaction between L-PRX and β4 integrin in myelinating Schwann cells. We conducted a biophysical characterization of the interaction using recombinant PRX-C and β4-FNIII-3.

Pulldown experiments with the purified components confirmed the direct molecular interaction suggested by the data above. His-tagged β4-FNIII-3 pulled down PRX-C (Figure 2A). Pulldown of a partially degraded PRX-C sample showed that essentially all PRX fragments on the SDS-PAGE gel were pulled down by β4-FNIII-3 (Figure 2B); MS analysis indicated that the shortest fragments were missing the very C-terminal regions, while the acidic domain was well-covered in all fragments (data not shown), suggesting that the N-terminal half of the PRX-C construct contained the binding site.

**Figure 2.**
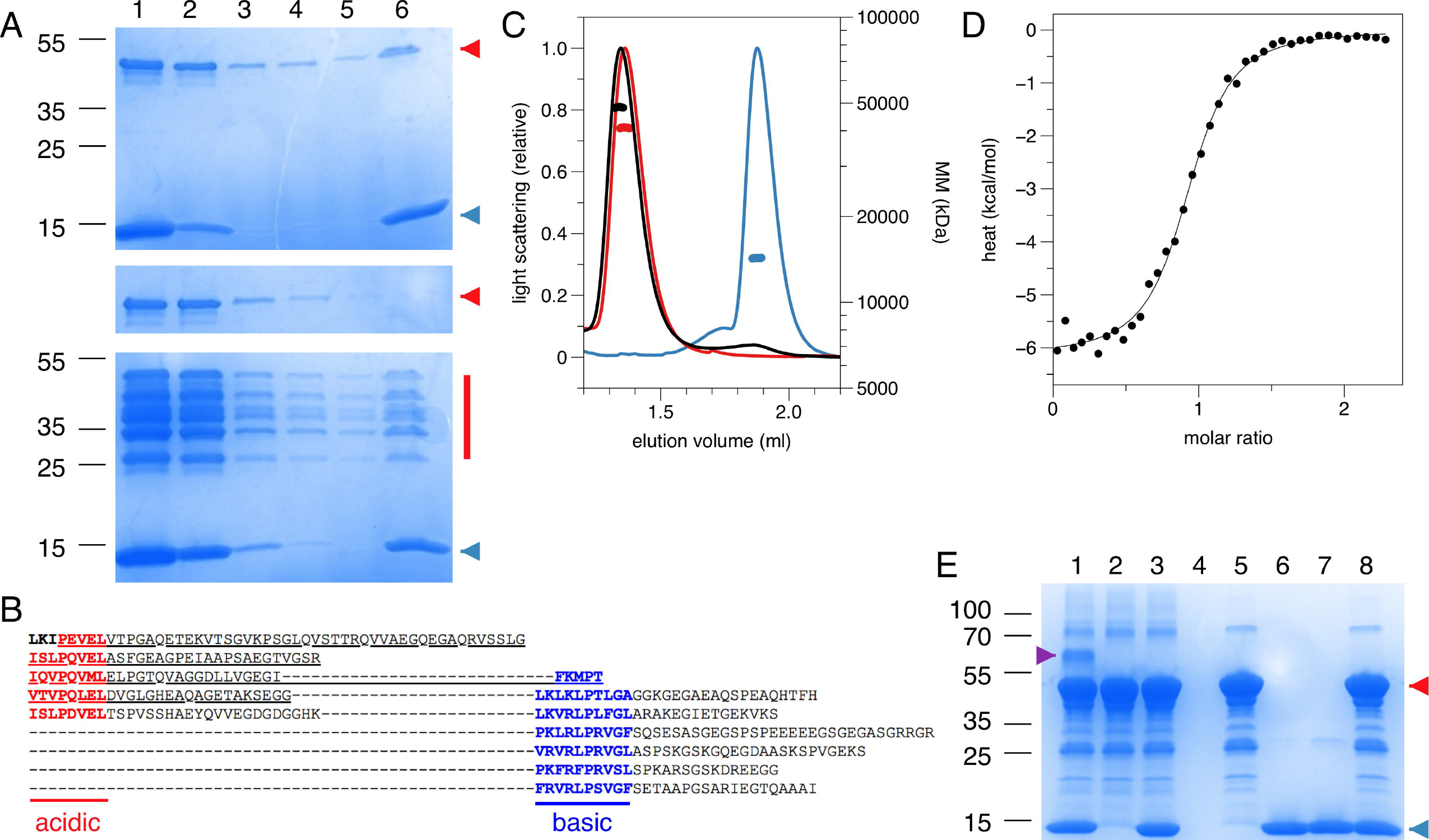
Direct molecular interaction assays using purified recombinant proteins. A. Pulldown with pure recombinant proteins on a Ni-NTA affinity matrix. Top, PRX-C and His-tagged β4-FNIII-3; middle, PRX-C alone; bottom; partially degraded PRX-C and His-tagged β4-FNIII-3. Samples: 1, input sample; 2, unbound fraction; 3-5, washes; 6, elution. PRX-C is indicated in red and β4-FNIII-3 in blue. Sizes of molecular weight markers are indicated on the left (kDa). B. Sequence of the PRX-C construct, indicating the presence of acidic and basic repeats. The underlined segment (the acidic domain) can be detected in all the PRX-C bands in the bottom panel of (A). C. SEC-MALS analysis shows molecular mass expected for a 1:1 complex. PRX-C, red; β4-FNIII-3, blue; complex, black. D. ITC titration of the complex indicates 1:1 stoichimetry. ΔH = 6.2±0.06 kcal/mol, ΔS = 5.8 cal/mol°. E. Covalent crosslinking. The complex is only visible, when both proteins have been activated (Lane 1). The magenta arrowhead indicates the 1:1 complex. PRX-C is indicated in red and β4-FNIII-3 in blue. Samples: 1, both proteins activated; 2, PRX-C activated; 3, PRX-C activated, β4-FNIII-3 added; 4, no activation; 5, no activation, PRX-C added; 6, no activation, β4-FNIII-3 added; 7, β4-FNIII-3 activated; 8, β4-FNIII-3 activated, PRX-C added.

SEC-MALS and QELS experiments verified the interaction *in vitro*, whereby the observed complex mass fit to a 1-to-1 complex, with an increased hydrodynamic radius (R_h_) compared to the individual interaction partners alone (Figure 2C, Table 1). R_h_ of the complex was only slightly higher than that for PRX-C, indicating that the complex remained in an extended overall conformation; this is also reflected in the very small change in SEC elution volume. As β4-FNIII-3 is small and tightly folded, this implicates that the PRX-C segment does not become compact upon complex formation, which would be indicated by a decreased R_h_.

**Table 1.**
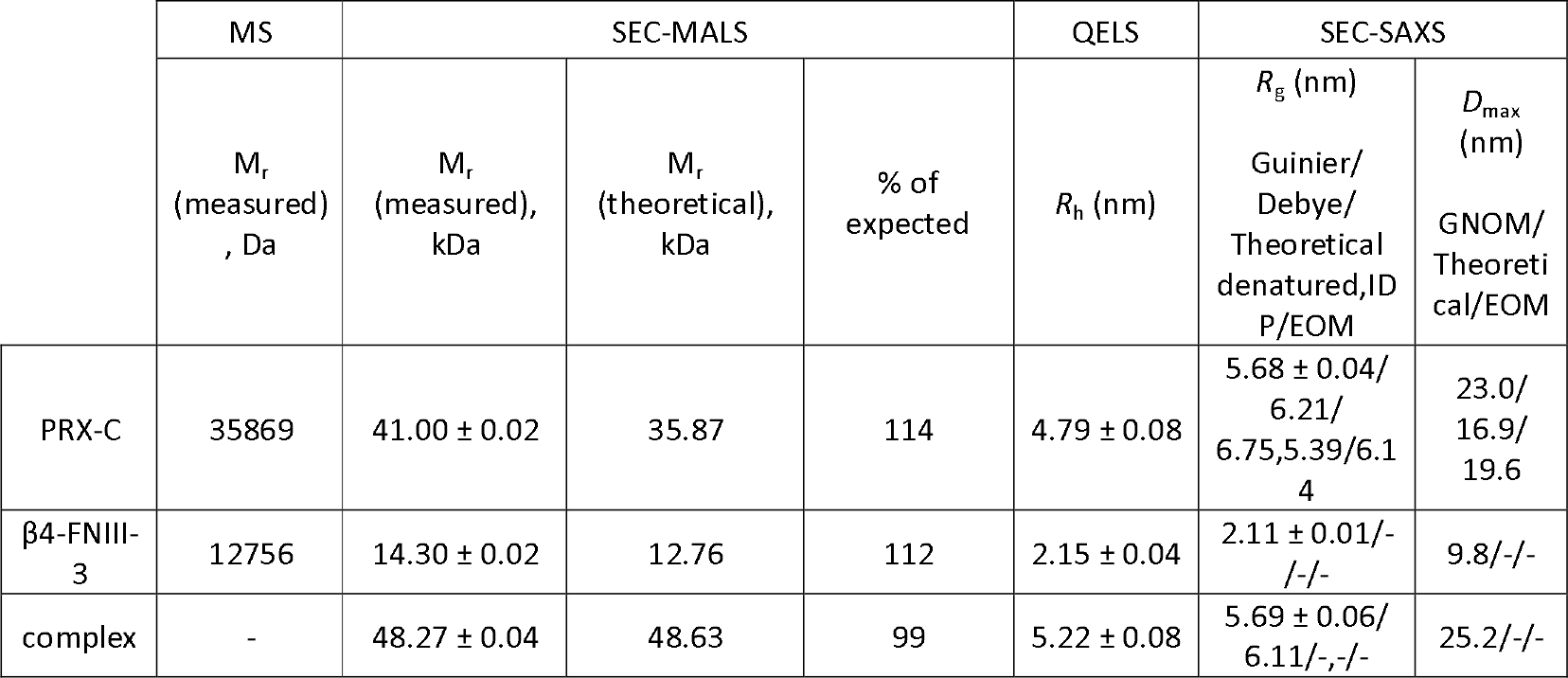
Characterization of the PRX-C/β4-FNIII-3 complex mass and size.

|  | MS | SEC-MALS |  |  | QELS | SEC-SAXS |  |
| --- | --- | --- | --- | --- | --- | --- | --- |
| | $M_r$<br>(measured)<br>, Da | $M_r$<br>(measured),<br>kDa | $M_r$<br>(theoretical),<br>kDa | % of<br>expected | $R_h$ (nm) | $R_g$ (nm)<br>Guinier/<br>Debye/<br>Theoretical<br>denatured, ID<br>P/EOM | $D_{max}$<br>(nm)<br>GNOM/<br>Theoretical/EOM |
| PRX-C | 35869 | 41.00 ± 0.02 | 35.87 | 114 | 4.79 ± 0.08 | 5.68 ± 0.04/<br>6.21/<br>6.75, 5.39/6.14 | 23.0/<br>16.9/<br>19.6 |
| β4-FNIII-3 | 12756 | 14.30 ± 0.02 | 12.76 | 112 | 2.15 ± 0.04 | 2.11 ± 0.01/-<br>/-/- | 9.8/-/- |
| complex | - | 48.27 ± 0.04 | 48.63 | 99 | 5.22 ± 0.08 | 5.69 ± 0.06/<br>6.11/-,-/- | 25.2/-/- |

A dissociation constant of 1.7 ± 0.1 μM was obtained for the protein-protein complex using ITC, with a binding stoichiometry of ∼1 (Figure 2D), being in good corroboration with the light scattering data above. Hence, the C-terminal region of L-PRX and β4-FNIII-3 bind each other with a low micromolar affinity, in a complex tight enough to survive *e.g.* separation by SEC. The stoichiometry suggests that the interaction site is either distinct from the repeat sequences in L-PRX, or that maximally one β4 integrin molecule can bind to PRX-C even if the binding site would involve the repeats.

Covalent crosslinking and mass spectrometry were used in an attempt to map the binding site in more detail. An additional band containing both proteins was apparent on SDS-PAGE after crosslinking (Figure 2E). MS analysis of tryptic peptides from this band confirmed the presence of both PRX-C and β4-FNIII-3.

Thermal stability of the proteins was studied using label-free differential scanning fluorimetry (nanoDSF), whereby the signal came from the folded β4-FNIII-3 domain. β4-FNIII-3 has the same melting point (+70°C) in the presence and absence of PRX-C (Table 2), suggesting no large structural changes are induced by PRX-C binding.

**Table 2.** Folding and thermal stability of PRX-C and β4-FNIII-3.

| | SRCD deconvolution* | | | Secondary structure prediction** | | | $T_m$, °C | |
| --- | --- | --- | --- | --- | --- | --- | --- | --- |
|  | % helix | % strand | % other | % helix | % strand | % other | SRCD | nanoDSF |
| PRX-C | 3<br>3 | 38<br>31 | 58<br>66 | 0 | 7.7 | 92.3 | - | - |
| β4-FNIII-3 | 1<br>2 | 45<br>50 | 51<br>49 | 0 (0) | 37.2 (34.5) | 62.8 (65.5) | 59.5 ± 0.6 | 69.8 ± 0.1 |
| complex | 2<br>3 | 39<br>35 | 56<br>61 | 0 | 14.9 | 85.1 | 58.8 ± 0.7 | 70.3 ± 0.1 |

\* Deconvolved using DichroWeb (values above), or with BeStSel (values below).
\*\* Predicted from primary structures using JPred. Values in brackets were extracted from the crystal structure using DSSP. The complex values were calculated from the JPred predictions, assuming a 1:1 complex.

### Structural insights into the protein-protein complex

While PRX-C is predicted to contain segments of β strand, the purified protein is essentially unfolded, as shown by SRCD spectroscopy (Figure 3A, Table 2). The β strand predictions coincide with repeats in the sequence (Figure 3B), and they could be of functional relevance in ligand protein binding and induction of secondary structures in PRX-C.

**Figure 3.**
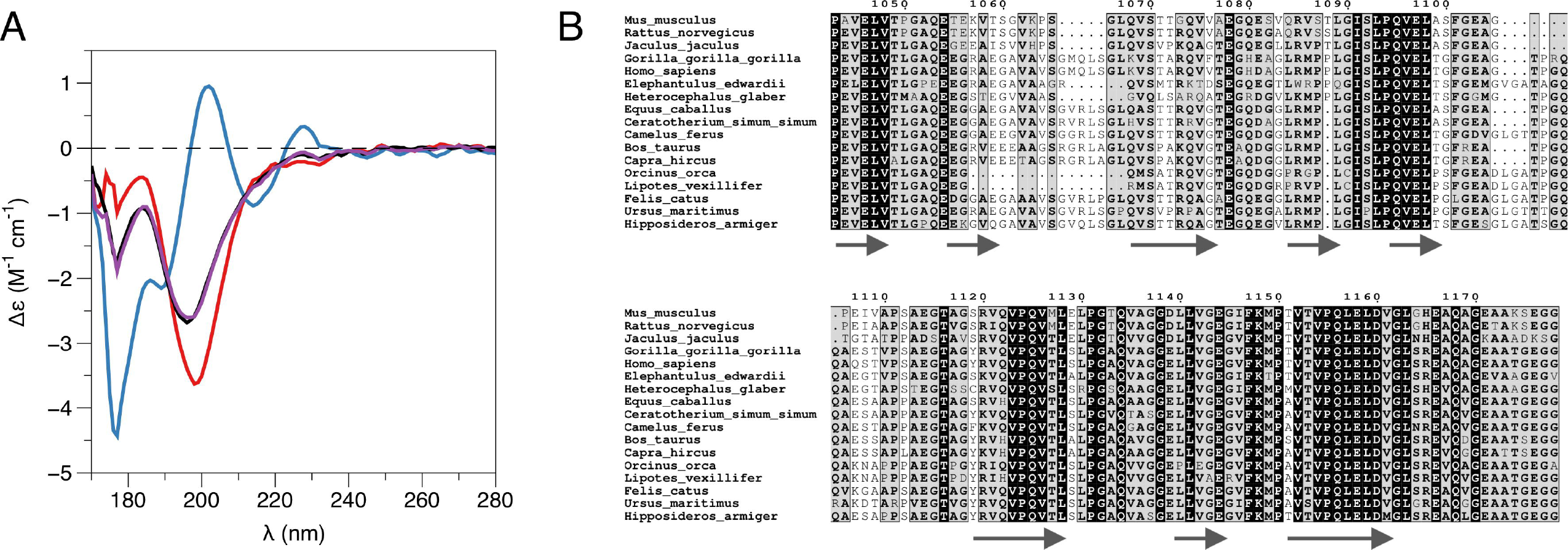
L-PRX remains disordered in the complex. A. SRCD spectroscopy. PRX-C, red; β4-FNIII-3, blue; complex, black; sum of individual protein spectra, magenta. B. Secondary structure predictions of the PRX-C acidic region, aligned from selected species.

SRCD was used to shed light on possible changes in secondary structure content accompanying complex formation. The spectra clearly show that the overall secondary structure content remains identical in the complex, compared to the two proteins in isolation (Figure 3A). Together with the SAXS data (see below), SRCD shows that upon β4-FNIII-3binding, the PRX C-terminal segment does not obtain large amounts of folded structure. Heat denaturation experiments by SRCD confirmed the same melting point for β4-FNIII-3 in the presence and absence of PRX-C (Table 2).

To aid in structural modelling and understanding the interactions in the complex, we solved the high-resolution crystal structure of the rat β4-FNIII-3 domain (Figure 4, Table 3). The structure is similar to the corresponding human protein (48), consisting of a β sandwich made of 7 β strands. In two of the four monomers in the asymmetric unit, the His tag could be partially seen, being in different conformations (Figure 4).

**Figure 4.**
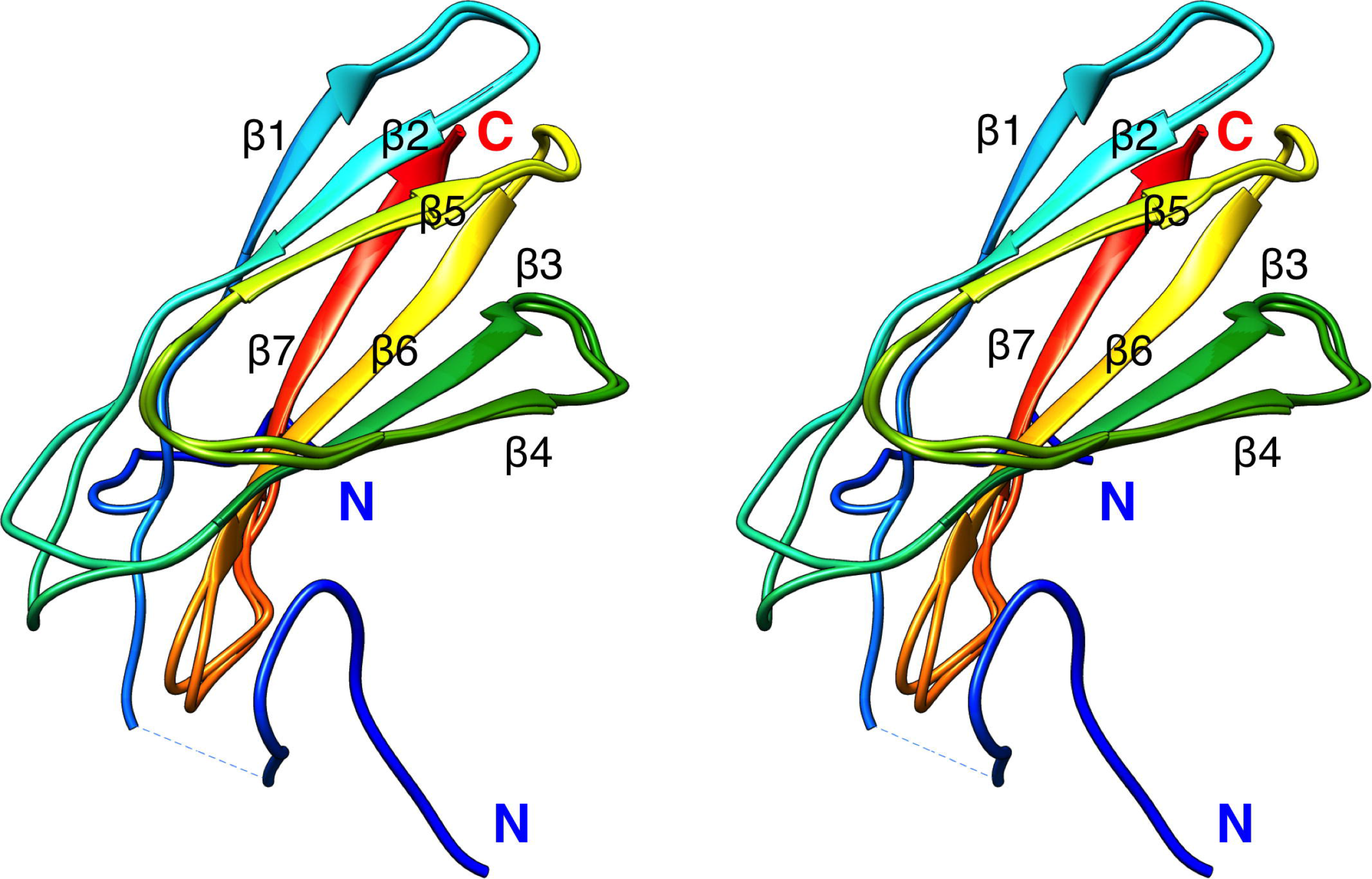
Crystal structure of rat β4-FNIII-3. Two of the 4 individual chains in the asymmetric unit are shown. The chain is coloured from the N (blue) to the C (red) terminus. The His tag is partially resolved at the N terminus, and takes different conformations in different monomers. Secondary structure elements are labelled.

**Table 3.** Crystallographic data collection and refinement statistics. Values in parentheses correspond to the highest-resolution shell.

|  |  |
| --- | --- |
| Data collection statistics |  |
| Beamline | ID30A-1 (ESRF) |
| Space group | P2<sub>1</sub>2<sub>1</sub>2<sub>1</sub> |
| Unit cell *a,b,c* (Å) | 51.22, 52.84, 144.53 |
| Wavelength (Å) | 0.966 |
| Resolution (Å) | 50-1.60 (1.64-1.60) |
| Unique reflections | 51757 (3746) |
| Multiplicity | 3.3 (3.3) |
| Completeness (%) | 98.3 (98.6) |
| R<sub>merge</sub> (%) | 6.1 (239.2) |
| R<sub>meas</sub> (%) | 7.3 (283.9) |
| <I/σ(I)> | 8.8 (0.6) |
| CC<sub>½</sub> (%) | 99.8 (16.6) |
| Wilson B (Å<sup>2</sup>) | 39.2 |
| Refinement statistics |  |
| R<sub>cryst</sub> (%) | 19.0 |
| R<sub>free</sub> (%) | 23.9 |
| RMSD bond length (Å) | 0.013 |
| RMSD bond angle (°) | 1.1 |
| Mean B value (Å<sup>2</sup>) | 49.7 |
| Ramachandran plot |  |
| Res. in favoured regions (%) | 95.9 |
| Outlier residues (%) | 0.8 |
| MolProbity score/percentile | 2.05 / 44<sup>th</sup> |
| PDB entry | 6HYF |

The purified PRX-C was subjected to SAXS experiments to gain more insight into flexibility, molecular dimensions, and conformational ensembles. These experiments indicate that PRX-C behaves much like a random polymer chain and is highly extended (Figure 5, Table 1).

**Figure 5.**
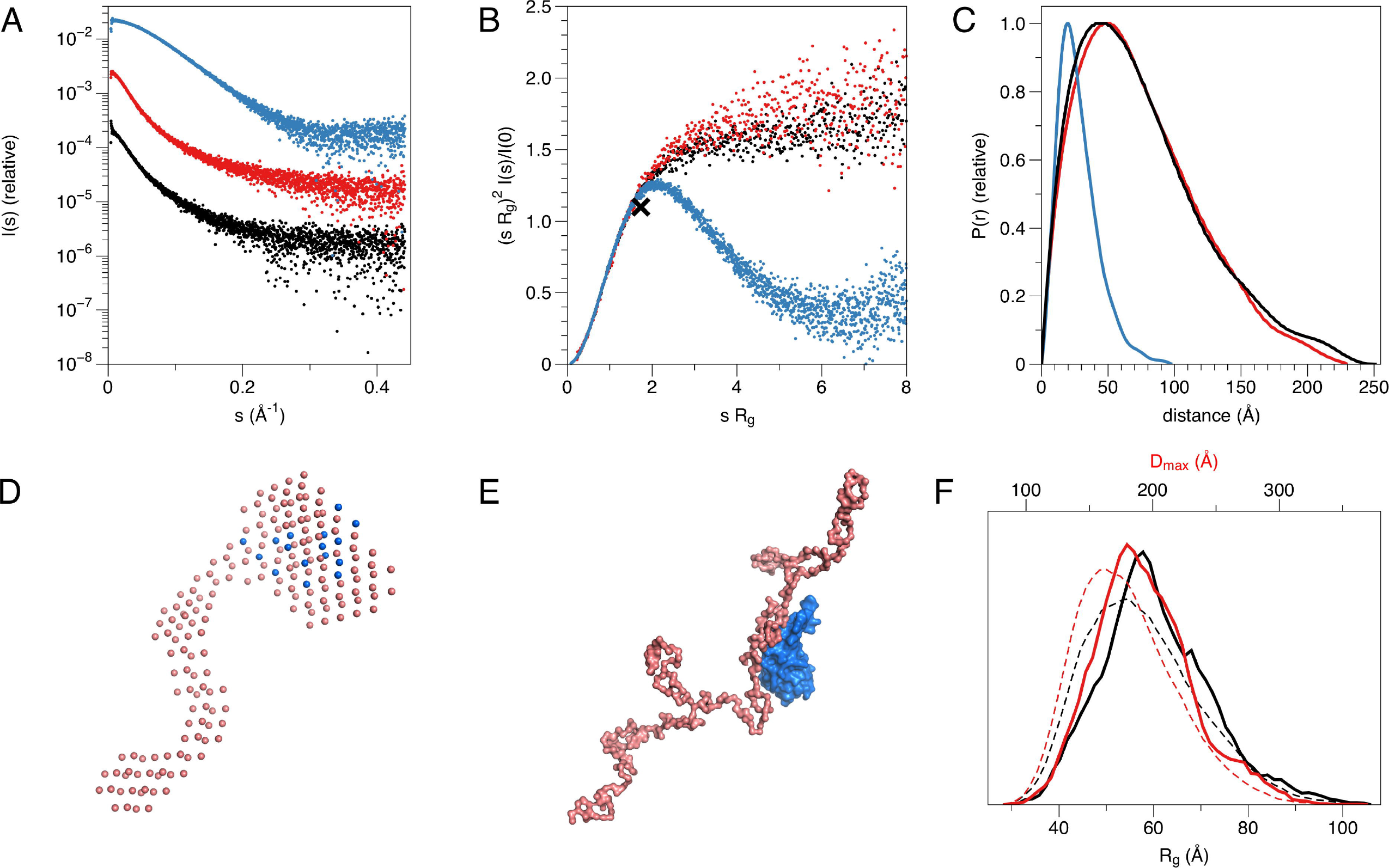
Structure of the complex in solution. A. SAXS scattering curve. PRX-C, red; β4-FNIII-3, blue; complex, black. B. Dimensionless Kratky plot. The black cross indicates the theoretical peak position for a folded, perfectly globular protein. C. Distance distribution function. D. Multi-phase model from MONSA. PRX-C, pink; β4-FNIII-3, blue. E. Rigid-body fit between a chain-like model of PRX-C (pink) and the integrin crystal structure (blue). F. EOM analysis for PRX-C. Dashed lines, theoretical distribution for a random coil; solid lines, the ensemble for PRX-C. Maximum dimension is shown in red and radius of gyration in black. PRX-C is slightly more elongated than a random coil.

While SAXS indicated a highly disordered nature for PRX-C alone, we wanted to see, whether it becomes more ordered in the complex with β4-FNIII-3. The β4-FNIII-3 crystal structure was used to model the solution structure of the complex, which cannot be crystallized due to the flexible nature of PRX and the fact that the exact binding site remains unknown. SAXS analysis of the protein complex, based on a SEC-SAXS experiment, shows that PRX-C remains elongated, and extra density corresponding to the size of β4-FNIII-3 appears close to one end of the complex (Figure 5D,E). This is in line with the results from SEC-MALS (see above).

The structure of β4-FNIII-3 can be used to predict the binding site for PRX. A surface analysis of the domain indicates an elongated hydrophobic groove lined by β strands 4 and 5 (Figure 4,6A), which could accommodate a linear motif, possibly in a β conformation; this could extend the smaller β sheet from 3 to 4 β strands. MD simulations of the structure further show that this region is the most mobile segment of β4-FNIII-3, and the cavity can open even more (Figure 6B-D). Furthermore, although the FNIII-3 and −4 domains of β4 integrin are rather homologous, sequence conservation in the possible binding site is very low - in line with the observation that FNIII-4 does not bind to PRX (Supplementary Figure 1).

**Figure 6.**
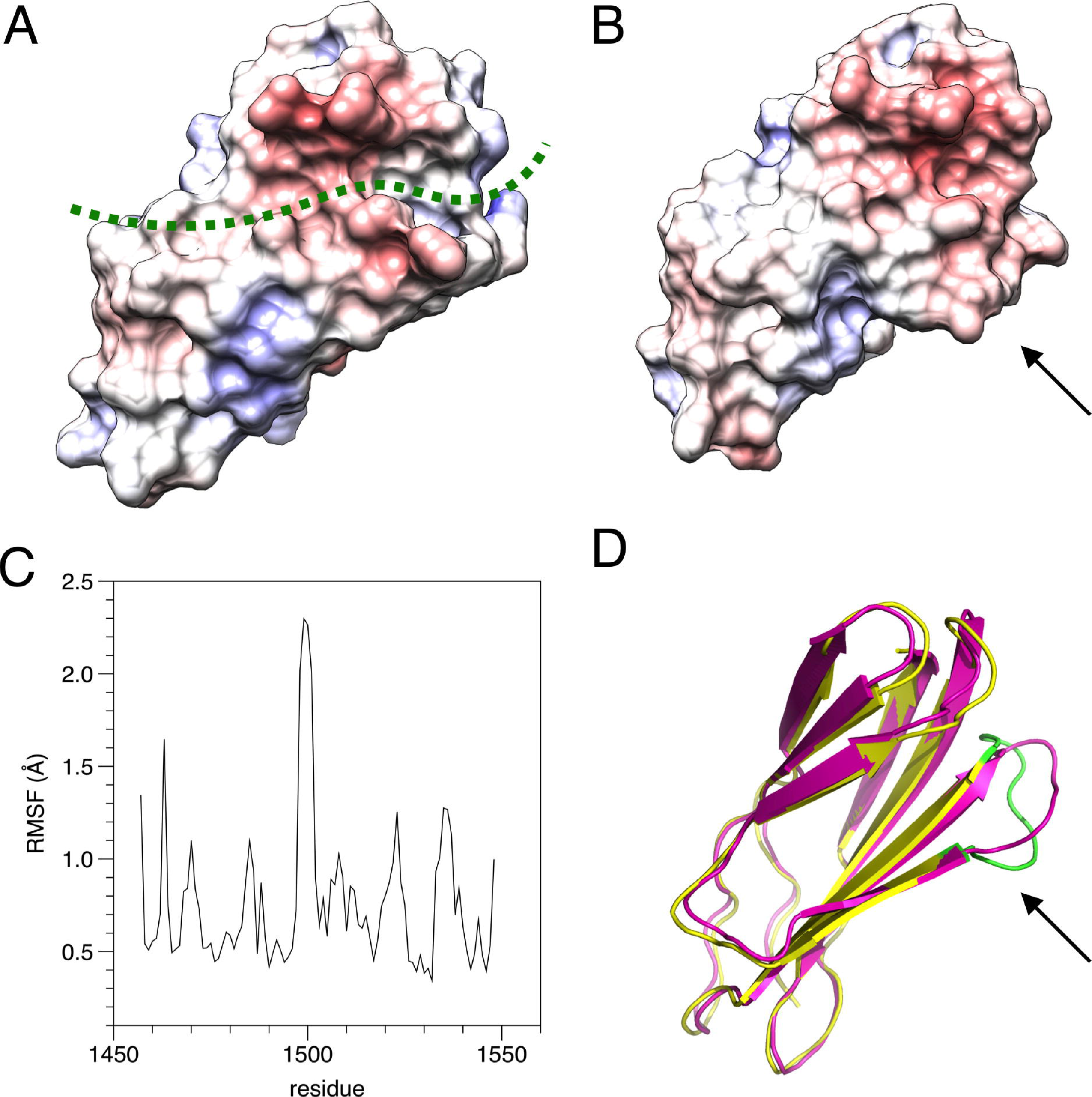
Possible binding site for L-PRX. A. Surface of the β4-FNIII-3 crystal structure, coloured by electrostatic potential. The predicted binding site is shown by a green dashed line. B. MD simulation indicates the possible binding site is flexible (arrow) and may open up more. C. RMSF during the 550-ns simulation. D. Comparison of the start (magenta) and end (yellow/green) points of the simulation. The loop at the potential binding site (arrow) is the most mobile region. The largest peak in the RMSF plot (panel C) is coloured green in the post-simulation structure.

## Discussion

At the cellular level, myelin formation involves a substantial amount of lipid and protein synthesis and their subsequent assembly into a multilayered, tightly packed proteolipid multilayer. The molecular interactions involved in this process can be roughly divided into those involved in the formation of the compact myelin compartment, and into those relevant for non-compact myelin. This division does not imply that a single protein cannot take part in both aspects; for example, MBP functions both in membrane packing and in the regulation of cytoplasmic channels.

Both PRX and β4 integrin play important roles in peripheral nerve myelination (11,15,25). Recently, it was suggested that PRX may be a ligand for β4 integrin (61), and the interaction has been highlighted before (16), but not studied in detail. We set out to confirm this putative direct molecular interaction, and to obtain 3D structural data on the protein-protein complex formed by PRX and β4 integrin.

### L-PRX binds β4 integrin directly with high affinity

Integrins are involved in myelination by both oligodendrocytes (CNS) and Schwann cells (PNS). In Schwann cells, the two main isoforms of integrin are α6β1 and α6β4. While α6β1 is expressed in the early stages of myelination and is crucial for the radial sorting of axons (62,63), α6β4 is predominant in the late stages and mature myelin (9,59). This indicates a developmental switch during myelination, and suggests β4 integrin is important for myelin maturation and maintenance. The levels of the Schwann cell laminin receptors integrin α6β1, α6β4 and dystroglycan are all regulated by the transcriptional co-activators Yap and Taz (63). The two basal lamina receptors of mature myelin, α6β4 integrin and dystroglycan, known to be together responsible for myelin stability (11), are according to our results bound to the same scaffold protein, L-PRX, in cytoplasmic membrane appositions of the Schwann cell abaxonal plasma membrane.

We have shown here with a collection of methods that L-PRX binds, through its C-terminal region, directly to β4-FNIII-3 with high affinity. Considering the presence of both proteins in a large protein scaffold at the Schwann cell outermost membrane, it is likely that the affinity is even higher, when full-length proteins interact and avidity is increased by *e.g.* protein clustering and oligomerization. PRX could function as a ruler between integrins and dystroglycans on the cell surface; on the other hand, changes in the clustering of these molecules, *e.g.* through changes in the basal lamina, might affect the conformation of PRX as well as its cytosolic interactions.

Similarly to the reduction seen here in the PRX interactome of β4 integrin in mice with truncated L-PRX, corresponding to human CMT4F mutation R1070X (24), a recent study (61) showed the loss of PRX from the β4 integrin interactome in mice with β4 integrin deficiency. It was thereby suggested that PRX might link β4 integrin functionally to PMP22 (61), but a direct or indirect interaction between PRX and PMP22 remains to be detected.

### L-PRX is intrinsically disordered

The C-terminal segment of L-PRX is highly flexible and intrinsically disordered, as shown by structure predictions, SRCD spectroscopy, and SAXS. The potential of intrinsically disordered regions in mediating specific protein interactions has been recently highlighted by a number studies (64–66). Such complexes involve usually the recognition of short linear motifs by a folded binding partner. In the current case, PRX-C remains highly flexible even upon β4 integrin binding. Considering the ratio R_g_/R_h_, the complex, having a lower ratio, is likely to be somewhat less flexible than PRX-C alone. Molecular flexibility may be important for the supramolecular organization of the protein scaffold at the outermost layer of PNS myelin, whereby PRX plays a central role.

Intriguingly, many of the myelin-specific proteins have a high degree of intrinsic disorder, as well as specific interactions, which makes myelin an interesting case for studying biomedically relevant intrinsically disordered proteins and their interactions. A well-studied example is the myelin basic protein, which is intrinsically disordered in solution, but partially folds upon membrane binding (67,68). MOBP and the cytoplasmic domain of P0 are predicteded to be disordered (69), and the latter has been experimentally shown to be intrinsically disordered in solution (70,71). The cytoplasmic domain of the large myelin-associated glycoprotein is an intrinsically disordered protein, but forms a specific heterotetramer upon dynein light chain binding, which is likely to help in dimerization and affect binding to the axonal surface (66).

In addition to the C-terminal segment studied here, most of the L-PRX-specific region is predicted to be intrinsically disordered. Hence, a monomer of L-PRX, assumed to be mainly in random coil conformation, could reach a length of >30 nm (72), which would be nearly doubled in the case of an L-PRX dimer formed through domain swapping at the N-terminal PDZ-like domain (22). PRX may thus be able to act as a molecular ruler and mediate protein interactions across partners distributed widely across the Schwann cell abaxonal membrane. The distance between bound DRP2 and β4 integrin can be estimated to be up to ∼25-30 nm, depending on the exact binding site and the conformation of L-PRX between the binding sites. The flexibility of L-PRX should allow it to stay bound to both protein complexes, even in the case they rearrange on the membrane.

### Insights from structure of the protein complex

The binding site for β4 integrin in L-PRX remains unknown, although the mapping in this study has suggested that the acidic region of L-PRX is involved. A sequence alignment of this region (Figure 3B) from different species shows that only a few segments are highly conserved; it is conceivable that these regions have functional relevance and they could form the binding site. Further studies will be required to obtain higher-resolution structural data to fully understand the binding interactions.

Typical modes of target protein binding by FNIII domains include transient opening and domain swapping (73). It is possible that some of the conserved segments in L-PRX are involved in a domain-swapping rearrangement of β4-FNIII-3. Such a mechanism would be compatible with the lack of change in secondary structure content upon complex formation, which we observed in SRCD experiments. High-resolution structural studies should answer many of the currently open questions, which will require the identification of the exact binding site(s).

Considering the structures of both L-PRX and β4 integrin, it is possible that additional binding surfaces are present in both proteins. β4 integrin has four FNIII domains in its cytoplasmic domain, and the C-terminal region of L-PRX is large and flexible, containing repetitive sequences. The fourth FNIII domain of β4 integrin does not bind to PRX-C (Supplementary Figure 1), however, and it is important to recapitulate that the stoichiometry of the interaction between PRX-C and β4-FNIII-3 was observed here to be 1:1.

### Relevance for understanding demyelinating disease

A number of proteins have been characterized as targets for mutations in human hereditary neuropathies, whereby the structure of the myelin sheath is compromised. Detailed knowledge about protein interactions in myelin is required to accurately understand the molecular bases of such diseases. One such protein is PRX, which is expressed as two isoforms, S- and L-PRX. While S-PRX consists of only a dimeric PDZ-like domain and a short tail, L-PRX dimerizes through the N terminus similarly to S-PRX (22), but has in total ∼1400 residues. Little is known about the structure and function of these long, repetitive L-PRX-specific segments. Expression of S-PRX may result in PRX heterodimerization (23) and affect the regulation of the PRX-containing protein scaffold, as S-PRX interacts with neither DRP2 nor β4 integrin.

Most PRX mutations causing human hereditary neuropathy introduce a premature stop codon into the sequence (74–76). In line with this, patients with such mutations have PRX expressed, but its size is smaller than in normal individuals (29,77). These observations hint at the possibility that an important function of L-PRX during myelin formation and maintenance lies at its C-terminal region. This is intriguing, given our observation that this region of L-PRX is intrinsically disordered. Here, we have shown a tight interaction between this C-terminal region and the third FNIII domain in the β4 integrin cytoplasmic domain – in the case of PRX truncations, such an interaction would be abolished. This is expected to cause larger-scale disturbances in the PRX-related protein scaffold reaching to the basal lamina and may translate to defects in myelin maintenance. Interactions with DRP2 are important for PRX function, but for a stable myelin sheath, links between the Schwann cell cytoplasm and the basal lamina through β4 integrin may be crucial. L-PRX, hence, mediates a two-pronged interaction from the cytoplasmic side through the Schwann cell plasma membrane to the basal lamina (Figure 7), and might have a role in sequestering different types of laminin receptors.

**Figure 7.**
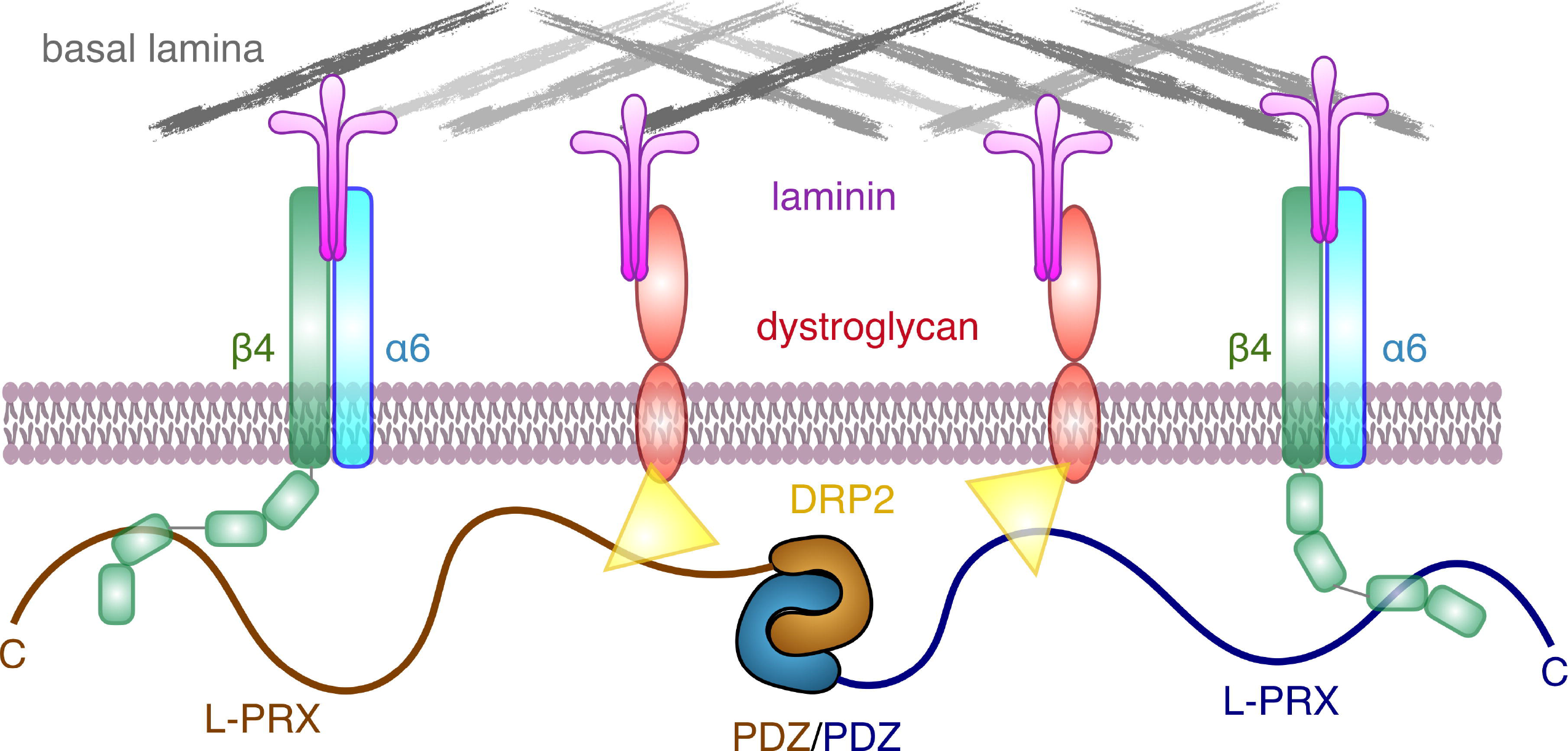
Schematic view of the PRX-linked protein scaffold in PNS myelin.

Loss of O-glycosylation causes similar phenotypic effects as L-PRX truncation in mouse models of CMT4F (78). Several O-glycosylation sites in PRX were identified, and some of them reside in the C-terminal region, which binds β4 integrin. It is possible that PTMs, including O-glycosylation, regulate PRX interactions with β4 integrin in a dynamic manner.

Integrin β4 has been highlighted both as a biomarker for Guillain-Barré syndrome (79–81), an autoimmune demyelinating disease of peripheral nerves, and as a central molecule in the entry of *Mycobacterium leprae* into Schwann cells, *via* the basal lamina (82). Leprosy is characterized by peripheral nerve damage initiated by mycobacterial infection of the Schwann cells. It is currently not known, how L-PRX binding to β4 integrin might modulate this process. Our data provide starting points for such studies in the future.

### Concluding remarks

The dimeric, highly elongated, flexible L-PRX is capable of acting as a central protein scaffold within non-compact myelin, linking integrins and dystroglycans together, thus bridging together two major protein complexes linking Schwann cells to the extracellular matrix. This could have high relevance in ensuring the necessary stability of membrane appositions that drive the formation of Cajal bands. The PRX-β4 integrin complex is likely to be important in both normal myelination, myelin maintenance, as well as the pathophysiology of neurodegenerative disease.

## Acknowledgements

We gratefully acknowledge the synchrotron radiation facilities and the beamline support at ASTRID2, DLS, ESRF, and EMBL/DESY. We express our gratitude towards the Biocenter Oulu Proteomics and Protein Analysis Core Facility and Dr. Ulrich Bergmann for providing access to mass spectrometric instrumentation, as well as Biophysics, Structural Biology, and Screening (BiSS) facilities at University of Bergen for providing calorimetric equipment and crystallization facilities. We thank Lisa Imrie for performing mass spectrometry and Qiushi Li for assistance. We are grateful to NanoTemper Technologies GmbH and Dr. Teresia Hallström for providing access to nanoDSF instrumentation.

## Author contributions

PB, DS, and PK conceived of the project. DS and PB carried out and analyzed yeast two-hybrid, immunochemical, and proteomics experiments. AR and HL performed experiments with purified proteins. AR and PK performed analysis of protein structure. AR, DS, and PK drafted the manuscript. All authors contributed to manuscript revision.

## Ethical statement

All animal work conformed to United Kingdom legislation (Scientific Procedures) Act 1986, and to the University of Edinburgh Ethical Review Committee policy.

## Data availability statement

Crystallographic data are available at the Protein Data Bank, under the accession code 6HYF. The raw data supporting the conclusions of this manuscript will be made available by the authors, without undue reservation, to any qualified researcher.

