## Supplementary Figure 1 for "Direct binding of the flexible C-terminal segment of periaxin to β4 integrin suggests a molecular basis for CMT4F"

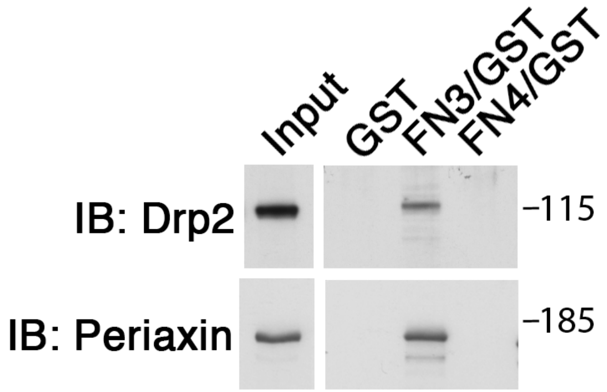


**Supplementary Figure 1. β4-FNIII-3 pulls down PRX and the associated DRP2 *in vitro*.** GST or a GST-β4-FNIII-3 (FN3/GST) fusion protein were incubated with a sciatic nerve lysate *in vitro* and bound L-PRX or DRP2 were detected by Western blotting (IB). GST-β4-FNIII-4 (FN4/GST) failed to pull down PRX and DRP2.
